# A Simple Parametric Representation of the Hodgkin-Huxley Model

**DOI:** 10.1101/2021.01.11.426189

**Authors:** Alejandro Rodríguez-Collado, Cristina Rueda

## Abstract

The Hodgkin-Huxley model, decades after its first presentation, is still a reference model in neuroscience as it has successfully reproduced the electrophysiological activity of many organisms. The primary signal in the model represents the membrane potential of a neuron. A parametric and simple representation of this signal is presented in this paper.

The new proposal is an adapted Frequency Modulated Möbius multicomponent model defined as a flexible decomposition in waves that describe the signal morphology. A specific feature of the new model is that the parameters are subject to interpretable restrictions.

A broad simulation experiment is conducted to show the new model accurately represents the simulated Hodgkin-Huxley signal. Moreover, the model potential to predict the neuron’s relevant characteristics, described with parameters of the Hodgkin Huxley model, is shown using different Machine Learning methods. The proposed model is also validated with real data from Squid Giant Axons. The comparison of the parameter configuration between the simulated and real data demonstrated the flexibility of the model as well as interesting differences.

**Author summary:** **Alejandro Rodríguez-Collado**. I received the double degree in Statistics and Computer Engineering and the Master’s degree in Business Intelligence and Big Data from the Universidad de Valladolid in 2019 and 2020, respectively. I work as researcher and Professor for the Department of Statistics and Operational Research at the Universidad de Valladolid. My main research interests include oscillatory signal processing, neuroscience, multivariate data analysis and supervised learning.

**Cristina Rueda**. I received the BS degree in mathematics from the Universidad de Valladolid in 1987 and the PhD degree in statistical science from the Universidad de Valladolid in 1989. I am currently Professor in the Department of Statistics and Operational Research at the Universidad de Valladolid. My main research interests include statistical inference methods under restrictions, circular data, computational biology, and statistical methods for signal analysis.

## Introduction

Neuroscience is an interdisciplinary science that studies the cellular, functional, behavioral, evolutionary, computational, molecular, and medical aspects of the nervous system. Many specialists from different areas of knowledge, such as physicists, chemists, mathematicians, computer engineers, and psychologists, have contributed to the field research. The mathematical approach is one of the most preferred ones, particularly in studying the electrophysiological activity between neurons. The signal that received most of the attention is the neuron membrane potential, which is the difference in electric charge between the cell’s interior and exterior. This signal is composed of various Action Potential Curves (APs). A single AP lasts a few milliseconds and consists of 3 stages: depolarization, repolarization, and hyperpolarization. For researchers, APs are of special importance: they are the informational unit between neurons, and their number and shape determine the morphological, functional, and genetic profile of the cell. For more detail see, [1], [2], [3], [4] or [5].

According to their dynamic behavior, neurons can be classified into excitable (generate an individual AP) or oscillatory (generate repetitive APs). Neurons that do not generate APs are called non-excitable. Neuronal cells of a similar type habitually exhibit identical behaviors. For instance, cardiac myocytes are usually oscillatory, while cortical neurons are mostly excitable. Furthermore, oscillatory neurons have been sub-classified in this work by the observed number of APs. A signal where several APs are observed is known as a Spike Train.

The most broadly considered mathematical model for describing AP dynamics is the Hodgkin-Huxley (HH) model, presented in [6]. More than a half-century later, this model remains key in neuroscience due to its innovative concept of modeling neuronal dynamics as a system of Ordinary Differential Equations (ODE) and its accurate representation of the electrophysiological neuronal activity. Specifically, the neuron’s membrane potential is stated to behave like an electrical circuit with three currents associated, each with three different types of ions. Sodium (Na^+^), potassium (K^+^) and another which is a non-specific leak current, mainly due to the influx of chlorine (Cl^−^). However, the HH model lacks identifiability as various parameter configurations can lead to the same observed signal. In addition, the model is not robust as minor manipulations of the values of the parameters can change its output completely ([7], [8]).

Many models that have been developed afterward are either simplifications or extensions of the HH model. Some emulate the HH model as biophysically realistic, whereas others seek more simple models. Among the first group are the Hopfield model and the Van der Pol oscillator’s extensions, such as the Fitz-Hugh Nagumo model. In the second group, some popular choices are the family of LIF models and the Izhikevich model. Some basic references about these models are [9], [10], [11], [12] and [13]. All the above are mechanistic models. The counterpart of the mechanistic models is the data-driven approach. Models based in data science, statistics and Machine Learning are in ever-rising popularity due to the increase in data availability and quality as noted in [14]. Our proposal can be framed within this class of models.

The Frequency Modulated Möbius (FMM) approach and others, such as the Fourier method, are encompassed in the AM-FM decompositions. A general overview of AM-FM decompositions and time-frequency signal analysis can be found in [15] and [16]. In particular, the FMM decomposition assumes a constant amplitude and a frequency that is modeled as a Möbius transformation. The monocomponent FMM model is presented in [17]. It shows how it accurately fits a wide variety of oscillatory patterns. The multicomponent FMM model is introduced in [18] and this concisely demonstrates its potential in neuroscience. Moreover, an exciting application for the automatic analysis of electrocardiograms is presented in [19].

This paper’s main goal is to show that the FMM model faithfully represents the AP signals derived from an HH model. To that end, a new FMM model is presented, denoted as FMM_ST_, where ST stands for Spike Train. It is an FMM multicomponent model with restrictions on the parameters. Expressly, the model assumes that the Spike Train is the concatenation of a fixed number of successive spikes with the same shape, each one described with a bi-component FMM.

In order to validate the model, a simulated experiment has been designed. A total of 5000 HH signals, corresponding to a wide variety of parametric configurations, has been generated according to a factorial design for the most relevant HH parameters. It is shown that the estimated FMM_ST_ signals accurately predict the simulated signal across all the parameter configurations. Furthermore, using Machine Learning, the potential of the FMM parameters to predict relevant HH parameters is also shown.

On the other hand, the model’s flexibility and good performance are also illustrated with the analysis of real data from the Squid Giant Axon Membrane Potential (SGAMP) database, originally from [20].

## Methods

### HH model

The presentation in this section follows those in [21] and [22]. The notation for HH variables and parameters is slightly changed from those used in these papers to avoid confusion with other terms introduced later in the paper.

The HH model is defined, see Definition 1 below, by a nonlinear ODE system for four state variables: the membrane potential (*X*) and the three opening probabilities of the ion gates (*m, n, h*). Furthermore, *X* depends on the input stimulus *I*(*t*) generated by other neurons’ post-synaptic currents. On their behalf, the variables *m, n* and *h* are referred to as voltage-gated channels as they depend on the membrane potential through the six ion flow rate functions denoted by *a_j_* and *b_j_*, *j* ∈ {*m, n, h*}, as explained in [4].

#### Definition 1.

Hodgkin-Huxley (HH) model.

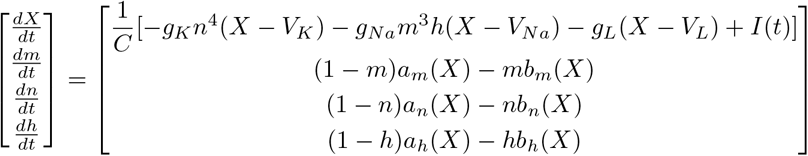

The first equation in Definition 1 depends on the parameters of the cell capacitance (*C*), the maximum conductances (*g_K_, g_Na_, g_L_*) and the equilibrium potentials (*V_K_, V_Na_, V_L_*) of the ionic currents. The six ion flow rate functions in rease the number of parameters to more than twenty parameters as explained in [22]. Hence, the parametric space of the model can be simplified by considering linear scalars of the ion flow rate functions 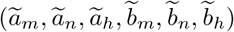, as done in [7] and [5].

An interesting simplification of the HH parametric space is the pair 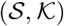 that has been recently considered in [7] and is defined as follows:

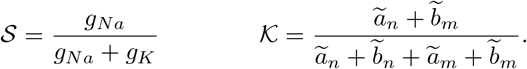

The authors of [7] explain the properties of the 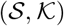 pair. In particular, they claim that the neuron’s excitability phenomenon is essentially bidimensional, being determined by the structure of the neuron as a cell and its ionic current kinetics. While 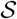 captures the neuron’s structural information, 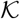 represents the kinetics of the ionic gates. Moreover, 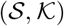 are less sensitive to slight changes in the signal than the primary HH parameters. However, such a drastic reduction in dimensions does not give a complete representation of the model.

### FMM approach

Let *t_i_*, *i* = 1,…,*n* denote the vector of observed time points and *X*(*t_i_*) the observed data, which in this paper is the potential difference in the neuron’s membrane. It is assumed that the time points are in [0, 2*π*). Otherwise, consider *t*’ ∈ [*t*_0_, *T* + *t*_0_] with *t*_0_ as the initial time value and *T* as the period. Transform the time points by 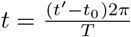.

Let, ***ν*** = (*A, α, β, ω*)’ be the four-dimensional parameters describing a monocomponent FMM signal, defined as the following *wave*: *W* (*t*, ***ν***) = *A* cos(*ϕ*(*t, α, β, ω*)), where *A* is amplitude and,

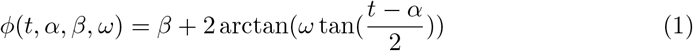

is the phase. Where, *α* is a location parameter and the parameters *β* and *ω* determine the skewness and kurtosis. More details about the parameters can be found in [17].

The FMM approach relies upon a signal plus error model, as follows:

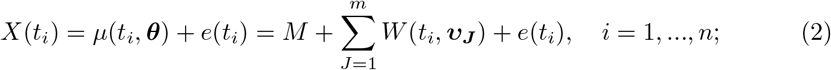

where, ***θ*** = (*M*, ***ν***_1_,…, ***ν*_*m*_**) and (*e*(*t*_1_),…,*e*(*t*_*n*_))’ ~ *N_n_*(0, *σ*^2^****I****).

The papers [19] and [18] consider particular FMM models, show the broad type of signals that the model represents, provide properties, and interpret the parameters as well as detail the algorithm used to fit the models. In particular, in the second paper, its potential in neuroscience is concisely shown.

Depending on the application, the waves of the model represent different physiological processes. For instance, in the ECG case, the waves of the FMM model represent the five fundamental ECG upward and downward deflections, which are universally named P, QRS complex (a wave complex), and T.

The AP signals are modeled using two (or three in some cases) components, the first being much more relevant than the rest. This component, denominated Dominant Component, identifies when the neuron spikes, allows the AP’s approximate reconstruction in the presence of noise, and the detection and identification of overlapping spikes. Moreover, theoretical properties are derived for the Dominant Component. All of these are shown in [18].

In Fig. 1, AP single signals and the FMM_ST_ predictions are shown. Fig. 1(A) illustrates the role of each parameter, while Fig. 1(B) shows the wave composition for an excitatory and an oscillatory neuron.

**Fig 1.**
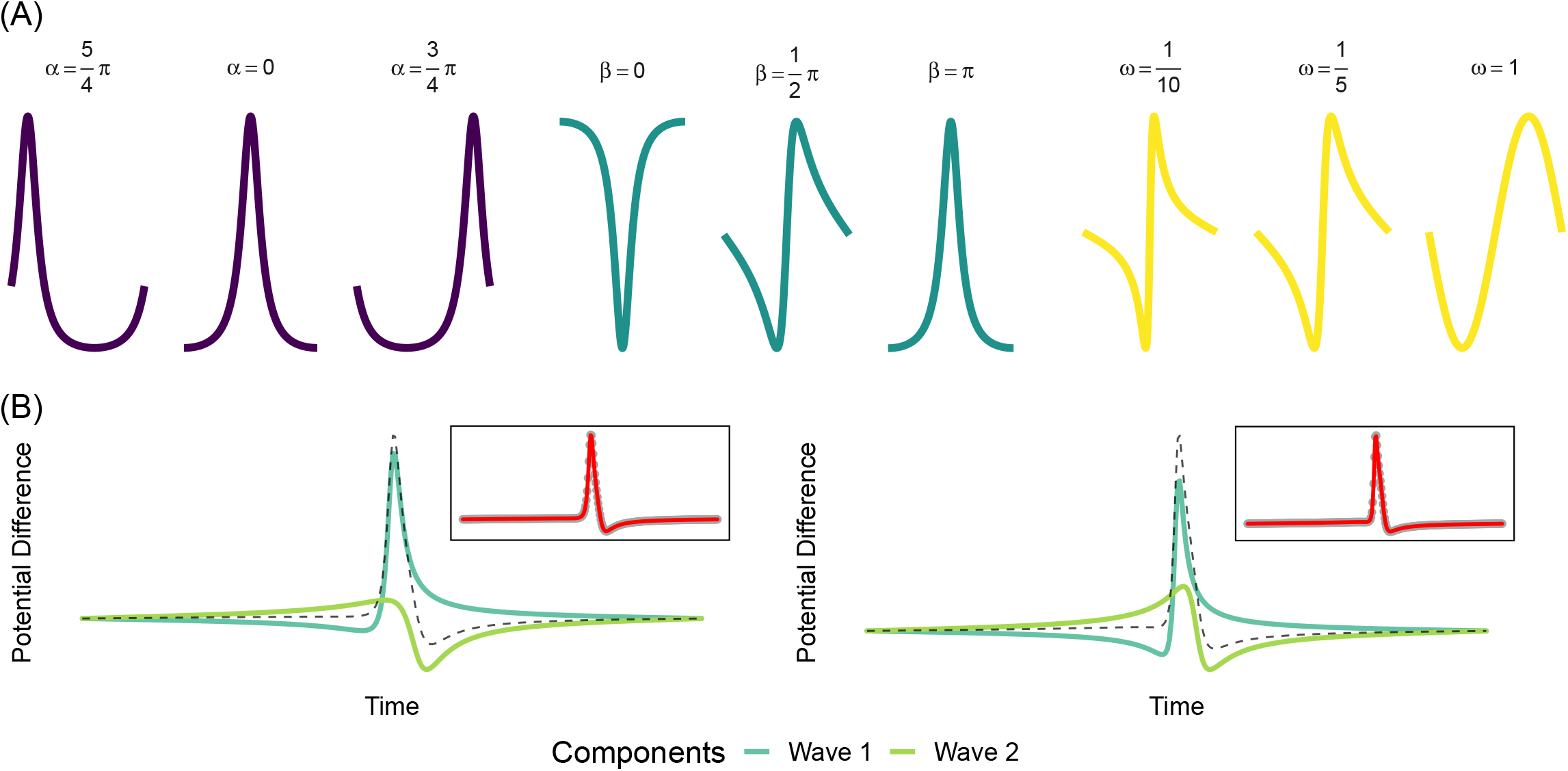
**(A)** Different FMM wave patterns generated by the parameters variation. **(B)** Components plots for the FMM_2_ models for an AP from an excitable neuron (left) and from an oscillatory neuron (right).

The FMM_ST_ model is a particular multicomponent FMM model, where restrictions on the parameters are imposed and is defined as follows:

#### Definition 2.

FMM_ST_ Model

For i=1,…,n:

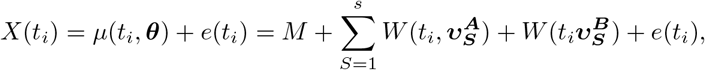

where,

- 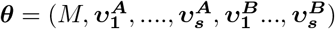
- 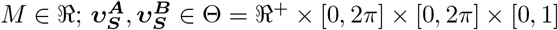; *S* = 1,…,*s*,
- 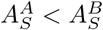; *S* = 1,…,*s*,
- 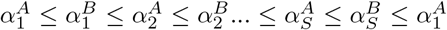
- 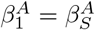 and 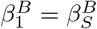; *S* = 2,…,*s*,
- 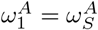 and 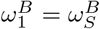; *S* = 2,…,*s*,
- (*e*(*t*_1_),…, *e*(*t_n_*))’ ~ *N_n_*(0, *σ*^2^***I***),

where *s* is the number of spikes that is assumed known and can be easily estimated with a naive method, as is detailed in the estimation algorithm section.

The model also assumes that each spike is modeled with two waves with parameters ***ν^A^*** and ***ν^B^***, respectively. The restrictions on the *A*s and the *α*s guarantee identifiability, and the restrictions on the *ω*s and *β*s represent the assumption of spikes having the same shape.

The number of free parameters of the FMM_ST_ is 1 + 4*s* + 4.

Additional restrictions may also be applied, depending on the application at hand. In particular, we will consider the following restrictions on the parameters *A* to analyse the SGAMP data:

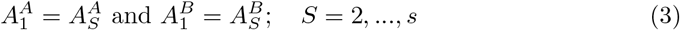

In this case, the number of free parameters is reduced to 1 + 2*s* + 6.

Moreover, other parameters of interest in practice are the distance between the *A* and *B* component for a given AP and the distance between consecutive APs. These are defined as follows,

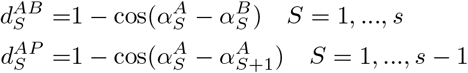

In some applications, it would be fair to assume 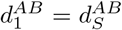; *S* = 2,…,*s*, which implies a reduction in the number of free parameters to 1 + *s* + 8.

It would also be suitable to assume that the distances between consecutive spikes are constant, for instance in controlled experiments without stimulus changes. The restrictions that apply for that case are:

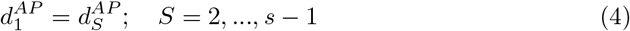

and the number of free parameters is then reduced to 1 + 1 + 8.

### Estimation algorithm

The implementation of FMM models in the programming language R, including applying the defined restrictions, is openly available in [23]. It is assumed that the segments to be analyzed represent complete spikes, in particular, *X*(*t*_1_)≃ *X*(*t_n_*). s is easily determined by a naive method based on a threshold as proposes [24]. This threshold is *k* = 2.5*σ_X_*, *σ_X_* being the sample standard deviation of the observed data.

Occasionally, two different parameter configurations represent a given signal equally well. However, one is physiologically more plausible. In that case, additional restrictions on the parameters are needed to guarantee that the solution is the one expected. In the case of SGAMP signals, it is assumed that a spike has a prominent Dominant Component and then that *A^A^* − *A^B^* > *C*, for a given threshold, *C*.

### Validation measure

The goodness of fit of the model is measured with an *R*^2^ statistic, which is the proportion of the variance explained by a model out of the total variance, as follows:

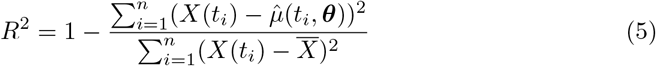

In this paper, 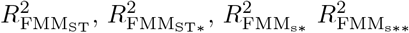 refer to the *R*^2^ value for the FMM_ST_ model, the FMM_ST_ model with restrictions on the *A*s, the FMM_*s*_ model with restrictions on *β*s and *ω*s, and the FMM_*s*\*_ model with restrictions on the *A*s, respectively.

### Machine Learning Supervised Methods

Several Machine Learning Supervised methods have been considered in the paper. At one end, the simple Linear Regression (LR) that serves as a benchmark approach. At the other extreme, the complex and “black box” Support Vector Machines of RBF Kernel (SVM) approach that has been proved to achieve excellent results in neuronal dynamics, as seen in [7] and [13], among others. Random Forest (RF) and Gradient Boosting Machines (GBM) are complex methods that provide interpretable results between the underfitting LR and the overfitting SVM, which have also been considered. [25] and [26] are essential references to learn about the procedures. The R packages [27], [28], [29] and [30], and the auxiliary package for learning procedures caret [31] have been used to implement the procedures.

### Programming languages

The simulation experiment has been developed combining the programming languages Python and R, which are probably the most used programming languages in data science. Python is used for data acquisition and transformation, while R fits the FMM models. Several solutions have been studied for the coupling between them that could, at the same time, be computationally effective, robust, and simple. A basic outline on the matter is presented in [32]. While certain libraries provide tools for the coupling of the two languages, such as [33], [34] and [35], these solutions are not sufficiently refined, and bash scripting was finally used.

## Results

### Simulation experiment design

In the first stage, Python simulates AP signals using a modified HH model implementation based on the one available in the Neurodynex package [21]. The original implementation offers several features, such as a detailed evolution of the voltage-gated variables *m, n, h*, or the application of input stimuli with different amplitude and shape. A modification was implemented to facilitate changes in the model parameters. The analyzed signal spans 60 ms. A short square input stimulus *I*, of just 1 ms, has been applied in the tenth ms of the simulation. See [36] to learn about the most used stimulus types. Five values for the strength of the stimulus have been selected, *I* = {0, 4.5, 7, 9.5, 12} *μA*, leading to a total of 5000 experiments. Different configurations of the most influential parameters have been considered according to a factorial experiment design. In every experiment, each of these parameters takes a random value from a set of preselected values. These values are experimentally observed in nature, as described in [7] and detailed in Table 1.

**Table 1.**
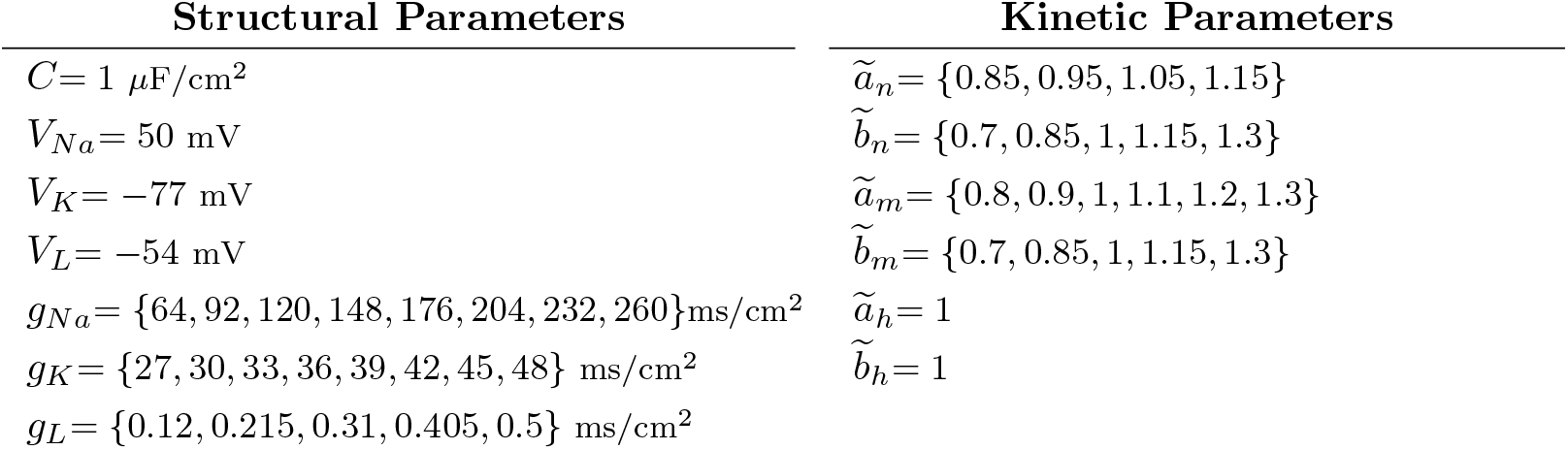
Parameters varied in the Hodgkin-Huxley simulations according to the designed factorial experiment design.

In a second stage, using R, the signals are preprocessed to assure that complete APs are analyzed. We proceed as follows: the first AP is discarded if it occurs before the input stimulus has been applied; resting potential values are considered instead of the discarded data to get segments of equal length for all the trials. An analogous procedure is applied if the last AP is observed in the last 6 ms of the experiment. In the exceptional case, where only a pre-stimulus AP is observed, the original signal is analyzed. It is relevant to note that discarding the first and/or last AP in the experiments is not a limitation of the approach, as we are assuming equally-shaped APs.

In a third step, an FMM_ST_ model is fitted to the data using the package [23].

### Simulation experiment results

3613 out of the total 5000 simulated HH experiments have been analyzed using the FMM approach as they have at least one AP. In Fig. 2, representative signals with one to four spikes are plotted along with the FMM_ST_ model predictions. There is a relevant Dominant Component that represents the depolarization and repolarization; and a second component that accounts for the hyperpolarization. A summary of the main statistics and parameter estimates are given in Table 2. The values in the table show the high prediction accuracy. In particular, the 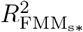 global mean (standard deviation) is equal to 0.8438 (0.0683), and the 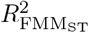 is equal to 0.9864 (0.0054). These values quantify what Fig. 2 shows. Table 2 also shows interesting differences in parameter configurations between signals with a different number of APs.

**Table 2.**
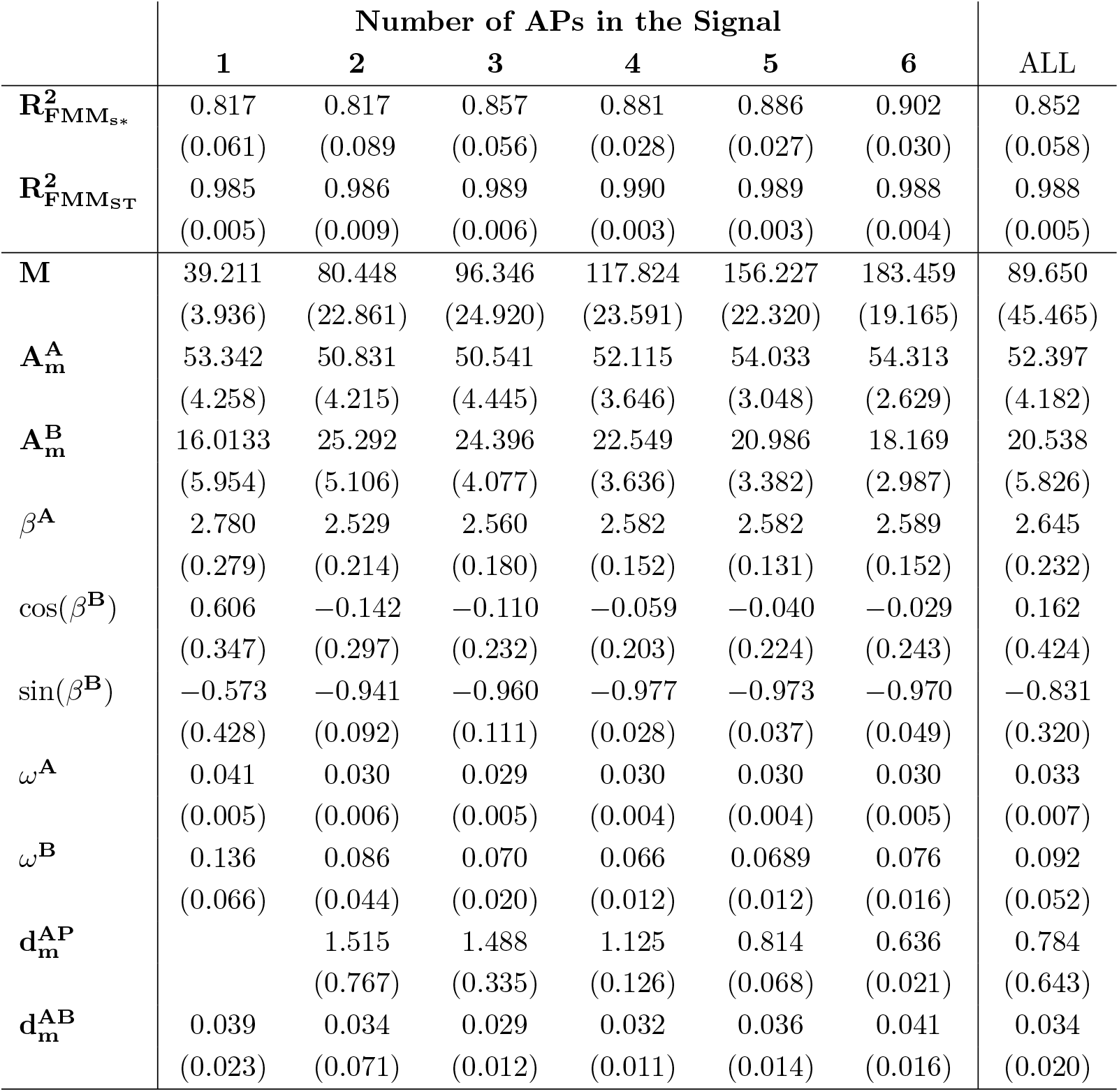
Means and standard deviations for *R*^2^ values and parameter estimators.

**Fig 2.**
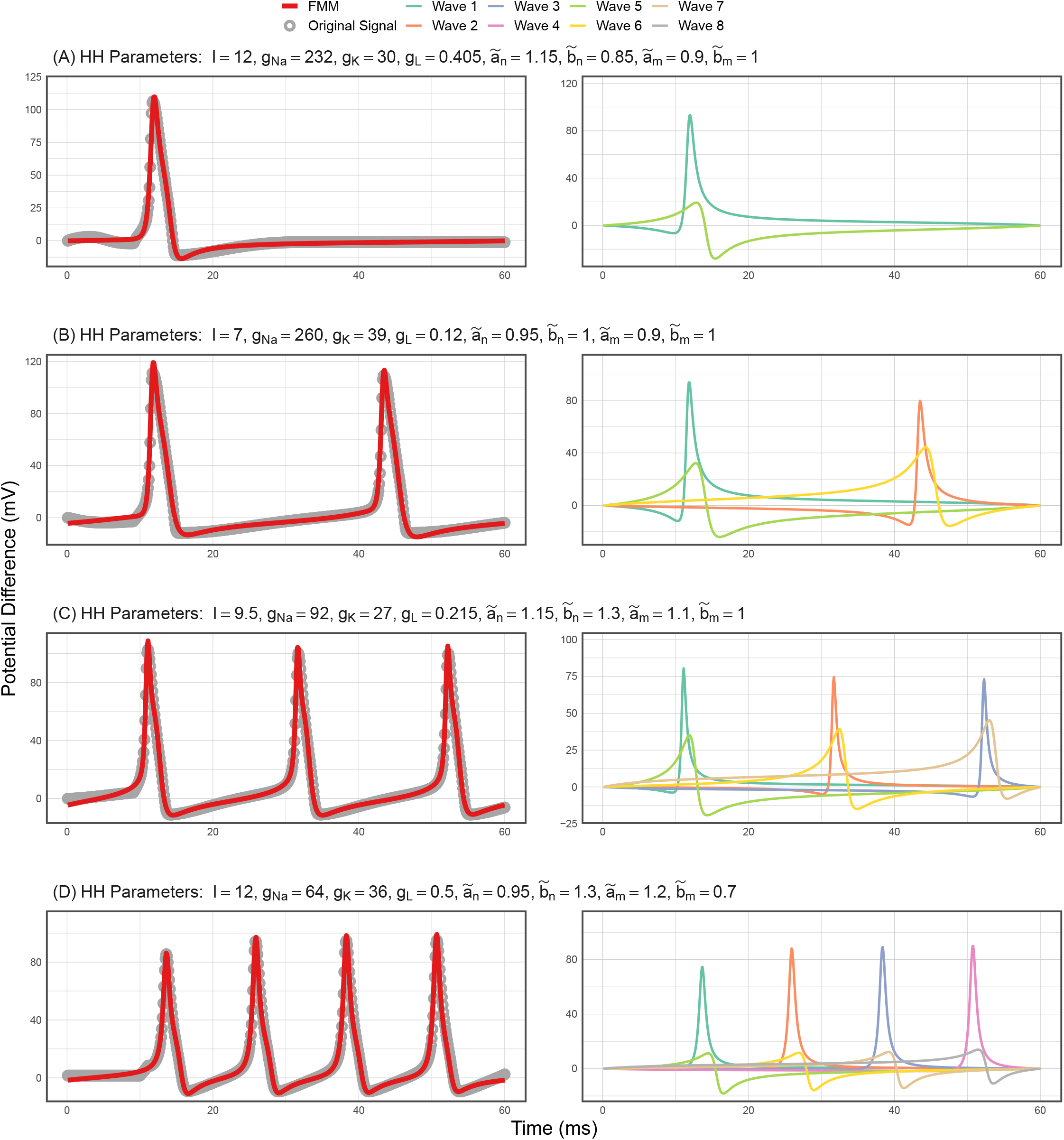
Neuronal signals simulated with the HH model and the estimated FMM_ST_ signals in red (left). Components of the fitted FMM_ST_ (right).

In particular, to illustrate how parameter configuration differentiates between excitable and oscillatory neurons in comparison with the HH parameters, two radar-plots corresponding to the HH and FMM models’ main parameters are represented in Fig. 3. The HH model parameters discriminate the two neuron types by biochemical properties. Specifically, the oscillatory experiments have primordially higher values in *g_Na_*, 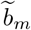 and 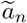. However, the FMM parameters differentiate the two types by the shape of the APs. It seems that excitable signals have both the components with a higher kurtosis and skewness (*ω^A^, ω^B^, β^A^, β^B^*) and a *B* component with a smaller amplitude 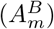.

**Fig 3.**
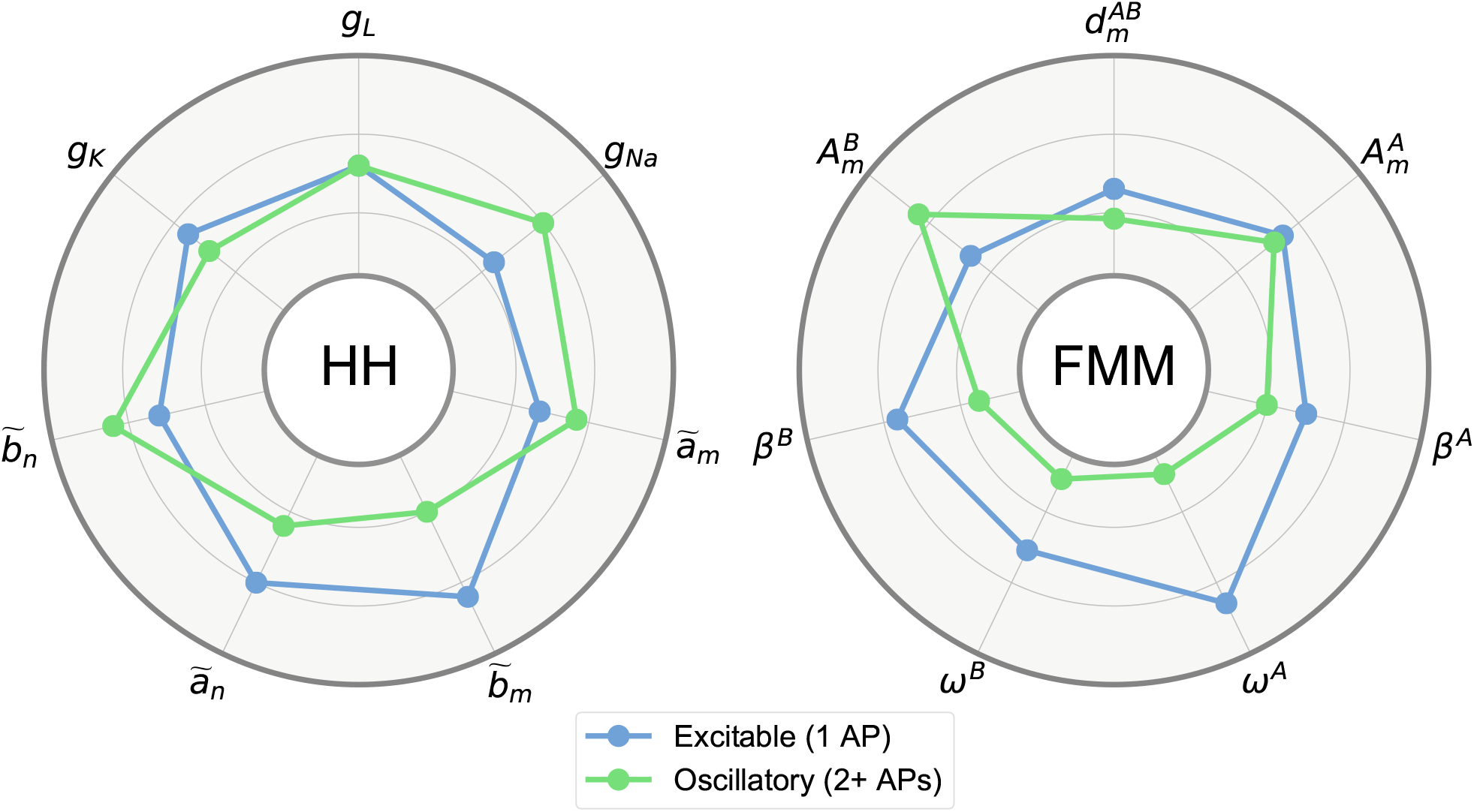
Median values of the HH parameters and FMM_ST_ parameters (defined in Table 3) by neuron type. The represented interval for each parameter is between the 10% and 90% percentiles.

Next, the analysis related to the 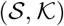 parameters is presented.

On the one hand, scatter plots of 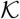 against 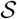 across intensities have been performed, imitating those in [7], in Fig. 4. Similar comments to those in the cited paper arise. Moreover, these plots also illustrate how the number of excitable and oscillatory experiments increase with the intensity. On the other hand, the potential of the model to predict the pair 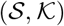 is shown. Two different sets of predictors: *τ^A^* and *τ^A^* + *τ^B^*, defined in Table 3 and different Supervised Learning procedures have been considered. Note that the LR and SVM approaches assume that the predictors are euclidean, but *β* is a circular parameter. Then, cos(*β^B^*) and sin(*β^B^*) are considered instead of *β^B^*. Furthermore, *β^A^* is considered as euclidean as it takes values concentrated in a small arc. Other predictors, derived from those in *τ ^A^* + *τ ^B^*, have been considered in the preliminary analysis but were eventually discarded using the principle of parsimony as only the results of LR improved lightly.

**Fig 4.**
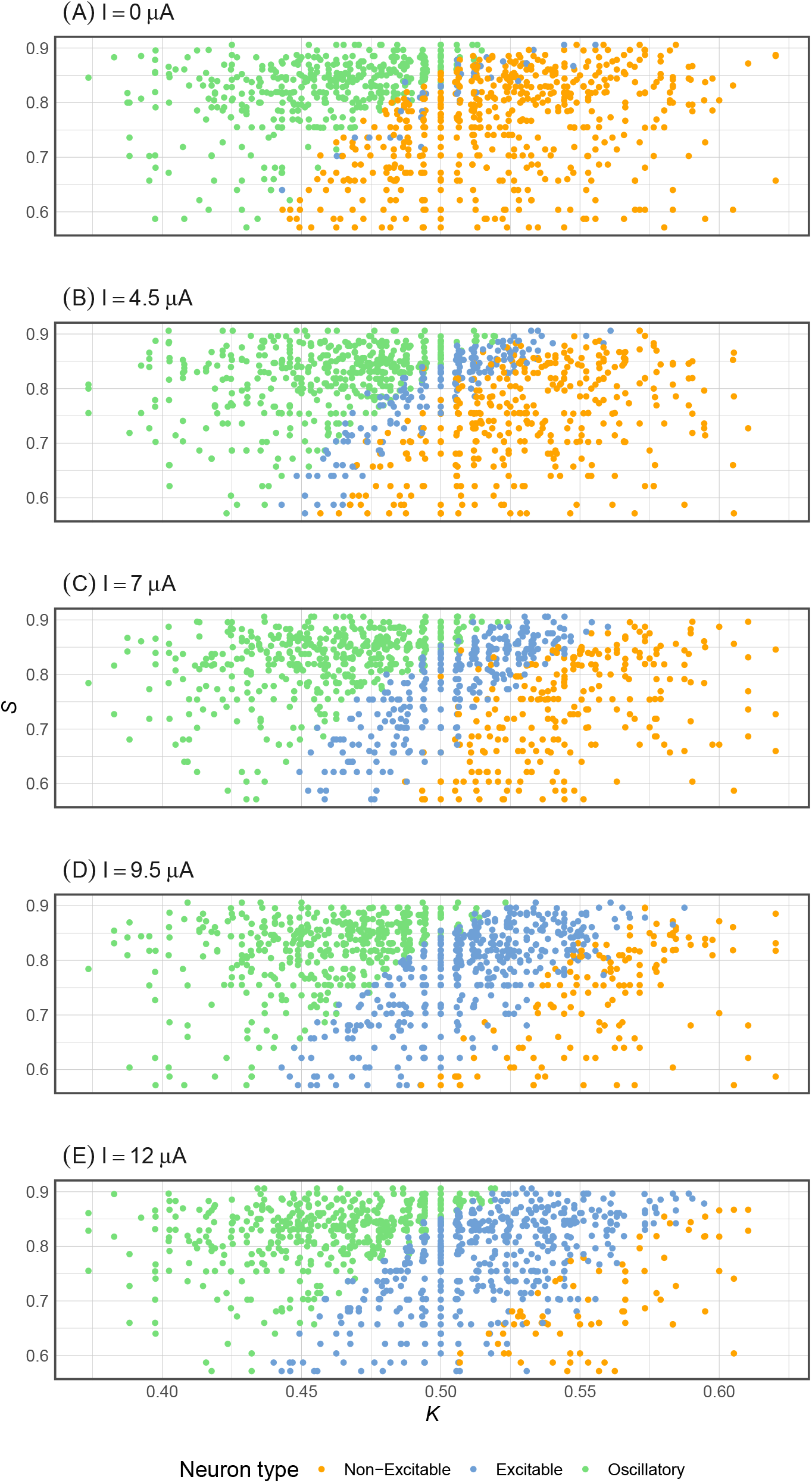
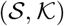 values by stimulus amplitude and neuron type.

**Table 3.** Feature sets used in the prediction of 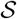 and 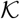.

| $\tau^A$ | | $\tau^B$ | |
| --- | --- | --- | --- |
| $M$ | Baseline level of the signal. | | |
| $A_m^A$ | Median of the amplitude $A_S^A$, $S = 1, \dots, s$. | $A_m^B$ | Median of the amplitude $A_S^B$, $S = 1, \dots, s$. |
| $\beta^A$ | Skewness of the $A$ waves. | $\beta^{B*}$ | Skewness of the $B$ waves. |
| $\omega^A$ | Kurtosis of the $A$ waves. | $\omega^B$ | Kurtosis of the $B$ waves. |
| $d_m^{AP}$ | Median of $d_S^{AP}$, $S = 1, \dots, s - 1$. | $d_m^{AB}$ | Median of $d_S^{AB}$, $S = 1, \dots, s$. |
| $s$ | Number of APs in the signal. | | |

\*: In linear procedures, $\cos(\beta^B)$ and $\sin(\beta^B)$ are used.

Variable selection methods have been used, a stepwise AIC for LR and caret’s RFE [31] for the rest. While in the former case, some variables are discarded, all the variables are included in the other models.

Furthermore, ten-fold cross-validation is considered for comparative and validation purposes. The dataset is divided into ten equally sized splits. In ten iterations, nine of the subsets are used to train the model, while the tenth serves as a test as in [25] and [37]. In addition, the Generalized Degrees of Freedom (GDFs) of the models have been calculated, as proposed in [38], to measure the underlying complexity.

Table 4 provides a summary of the results for the 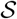 and 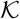 models. It can be seen that the parameters associated with the second component, *τ^B^*, significantly increase the prediction accuracy compared with that obtained using only parameters associated to component *A*. Regarding the different procedures, at one end, the relatively bad results for LR provides evidence that the relation between the predictors and 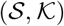 is not linear. At the other end, SVM gives the most accurate prediction, with more than 95% and 94% of explained variance for 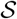 and 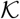 respectively. The RF and GBM results are between those for LR and SVM; while RF is comparable to GBM in terms of interpretability and complexity, the attained accuracy is less. However, although GBM procedures remain more complex and slightly less accurate than SVM, the predictions are interpretable. Specifically, the most relevant predictors to explain 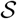 are *β^A^* and *ω^A^*, which implies that 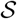 is well predicted by the AP shape. In the case of 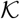, the most relevant predictors are *M*, 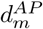 and *s*.

**Table 4.**
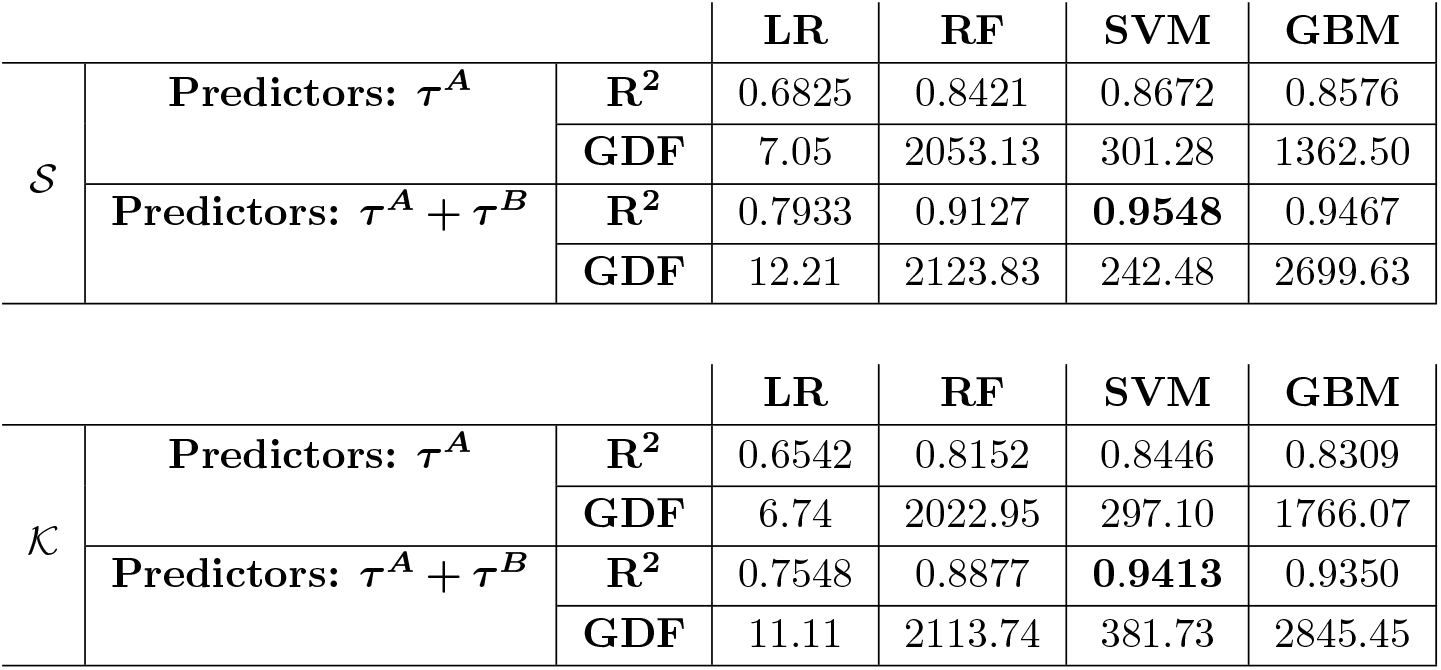
*R*^2^ and GDFs for the LR, RF, SVM and GBM models to predict 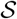 and 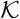.

|  |  |  | LR | RF | SVM | GBM |
| --- | --- | --- | --- | --- | --- | --- |
| S | Predictors: $\tau^A$ | $R^2$ | 0.6825 | 0.8421 | 0.8672 | 0.8576 |
|  |  | GDF | 7.05 | 2053.13 | 301.28 | 1362.50 |
| | Predictors: $\tau^A + \tau^B$ | $R^2$ | 0.7933 | 0.9127 | **0.9548** | 0.9467 |
|  |  | GDF | 12.21 | 2123.83 | 242.48 | 2699.63 |
|  |  |  | LR | RF | SVM | GBM |
| K | Predictors: $\tau^A$ | $R^2$ | 0.6542 | 0.8152 | 0.8446 | 0.8309 |
|  |  | GDF | 6.74 | 2022.95 | 297.10 | 1766.07 |
| | Predictors: $\tau^A + \tau^B$ | $R^2$ | 0.7548 | 0.8877 | **0.9413** | 0.9350 |
|  |  | GDF | 11.11 | 2113.74 | 381.73 | 2845.45 |

Predictive analysis for other HH parameters has also been performed. The results for sodium parameters are included in Table 5. Sodium parameters have been selected as they are more accurately predicted and are the engine of the neuron excitability, as some authors such as [39] and [8] claim. It is interesting to note that the 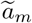 and 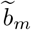 are directly related to the first and second FMM components, respectively, while *g_Na_* is related to both. The models’ accuracy is not as high as for 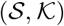, which are more stable parameters.

**Table 5.**
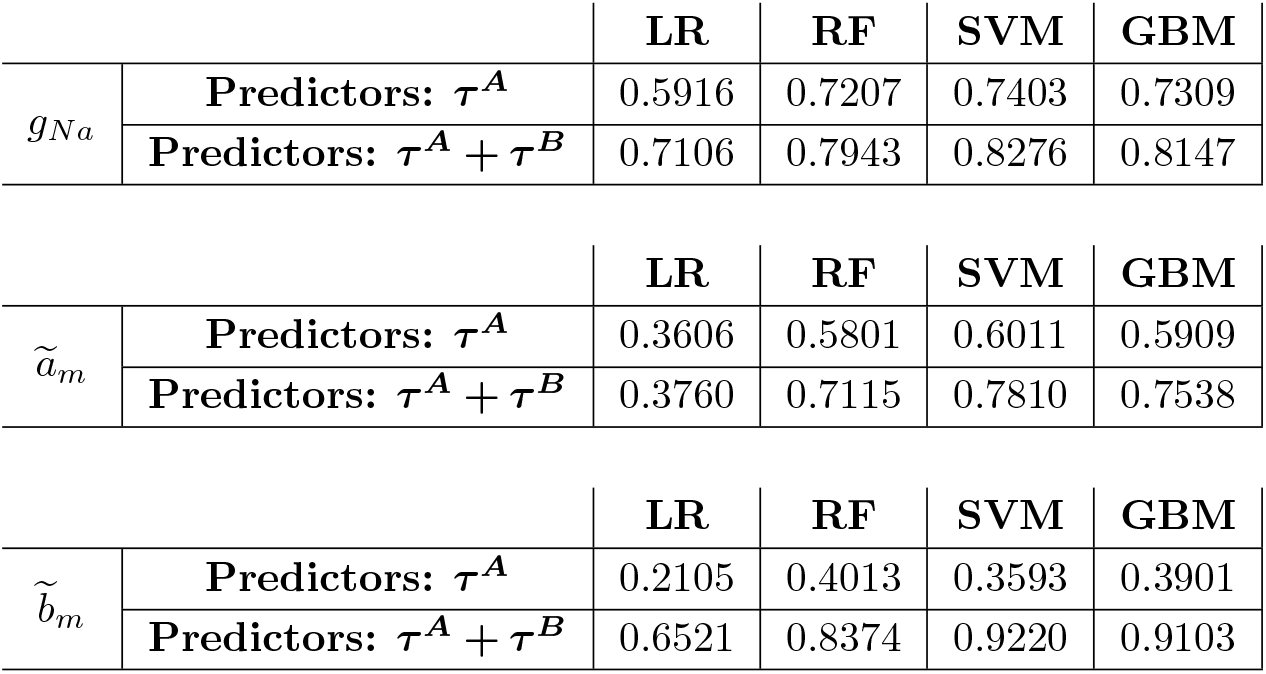
*R*^2^ values for LR, RF, SVM and GBM models to predict sodium parameters.

|  |  | LR | RF | SVM | GBM |
| --- | --- | --- | --- | --- | --- |
| $g_{Na}$ | **Predictors: $\tau^A$** | 0.5916 | 0.7207 | 0.7403 | 0.7309 |
|  | **Predictors: $\tau^A + \tau^B$** | 0.7106 | 0.7943 | 0.8276 | 0.8147 |
| $\tilde{a}_m$ | **Predictors: $\tau^A$** | 0.3606 | 0.5801 | 0.6011 | 0.5909 |
|  | **Predictors: $\tau^A + \tau^B$** | 0.3760 | 0.7115 | 0.7810 | 0.7538 |
| $\tilde{b}_m$ | **Predictors: $\tau^A$** | 0.2105 | 0.4013 | 0.3593 | 0.3901 |
|  | **Predictors: $\tau^A + \tau^B$** | 0.6521 | 0.8374 | 0.9220 | 0.9103 |

### Real Data

The SGAMP database, firstly used in [20] and publicly accessible at [40], contains single-unit neuronal recordings of North Atlantic squid (*Loligo pealei*) giant axons in response to stimulus currents. The database has been extensively used in works. Among the recent ones are [41] and [42]. The experiments where the applied stimulus is a short square stimulus, the same as in the HH experimentation, have been selected for the analysis. Five signals have been extracted from 4 trials of 3 different axons the stimulus amplitude being equal to *I* = 5 *μA* in the five cases. The length of the analyzed segment is also equal to that of the simulated HH signals, 60 ms, to facilitate comparisons.

Four different FMM models have been fitted to the signals: an FMM_s**_, FMM_s*_, FMM_ST_ and an FMM_ST*_. Moreover, it is assumed that *A^A^ – A^B^* < *C*, where *C* is 0.70 times the maximum difference obtained in previous iterations of the algorithm. For comparative proposes, Fourier models with the same number of free parameters as the FMM models, denoted as FD_*a*_ where *a* is the number of harmonics, have also been fitted.

Fig. 5 shows the FMM_ST*_ and the corresponding Fourier predictions for a representative signal. The *R*^2^ values and the number of free parameters are given in Table 6, for the five signals and eight models. The figures in the table show that the Dominant Component explains much of the variability here, much more than it does in the HH experimentation. Furthermore, the models with restrictions in the *A* parameters are as accurate as the others without them. As such, the former models, being the simpler ones, are preferred due to the parsimony principle. Besides, the FD_*a*_ models make much less accurate predictions. Moreover, Table 7 gives the parameter values for the FMM_ST*_ model of the five signals. Compared to the values in table 2, in particular to those corresponding to signals with 4 APs, some interesting differences can be highlighted, in particular in *A^A^*, *A^B^*, *β^A^*, and *β^B^*.

**Table 6.** *R*^2^ values of the different FMM and FD models for the signals extracted from the SGAMP database.

|  | N<sup>o</sup> of parameters | Experiment IDs |  |  |  |  |
| --- | --- | --- | --- | --- | --- | --- |
|  |  | a1t18_1 | a1t18_2 | a1t22 | a3t03 | a4t13 |
| **FMM<sub>S\*\*</sub>** | 8 | 0.9435 | 0.9457 | 0.9472 | 0.9512 | 0.9324 |
| **FMM<sub>S\*</sub>** | 11 | 0.9485 | 0.9511 | 0.9476 | 0.9566 | 0.9429 |
| **FMM<sub>ST\*</sub>** | 15 | 0.9862 | 0.9848 | 0.9880 | 0.9936 | 0.9910 |
| **FMM<sub>ST</sub>** | 21 | 0.9865 | 0.9852 | 0.9882 | 0.9937 | 0.9910 |
| **FD<sub>4</sub>** | 9 | 0.3635 | 0.3609 | 0.3587 | 0.4080 | 0.2809 |
| **FD<sub>5</sub>** | 11 | 0.3648 | 0.3622 | 0.3594 | 0.4141 | 0.2865 |
| **FD<sub>7</sub>** | 15 | 0.3687 | 0.3636 | 0.3614 | 0.4305 | 0.3056 |
| **FD<sub>10</sub>** | 21 | 0.5676 | 0.5809 | 0.5586 | 0.6099 | 0.4695 |

**Table 7.** Parameter estimators of the FMM_ST*_ models for the SGAMP signals.

|  | Experiment IDs |  |  |  |  | Mean |
| --- | --- | --- | --- | --- | --- | --- |
|  | a1t18_1 | a1t18_2 | a1t22 | a3t03 | a4t13 |  |
| **M** | 173.617 | 179.144 | 115.622 | 158.667 | 184.105 | 162.231 |
| **A<sup>A</sup>** | 44.955 | 44.472 | 37.352 | 43.089 | 46.284 | 43.230 |
| **A<sup>B</sup>** | 6.893 | 6.770 | 13.989 | 6.478 | 27.524 | 12.331 |
| **$\beta^A$** | 3.426 | 3.311 | 3.933 | 3.572 | 3.634 | 3.575 |
| **$\cos(\beta^B)$** | -0.368 | -0.387 | -0.214 | -0.428 | -0.221 | -0.324 |
| **$\sin(\beta^B)$** | -0.930 | -0.922 | 0.977 | -0.903 | 0.975 | -0.161 |
| **$\omega^A$** | 0.028 | 0.028 | 0.030 | 0.027 | 0.027 | 0.028 |
| **$\omega^B$** | 0.101 | 0.138 | 0.008 | 0.097 | 0.005 | 0.070 |
| **$d_m^{\text{AP}}$** | 1.016 | 0.985 | 0.987 | 1.033 | 1.049 | 1.014 |
| **$d_m^{\text{AB}}$** | 0.021 | 0.020 | 0.002 | 0.020 | 0.002 | 0.013 |

**Fig 5.**
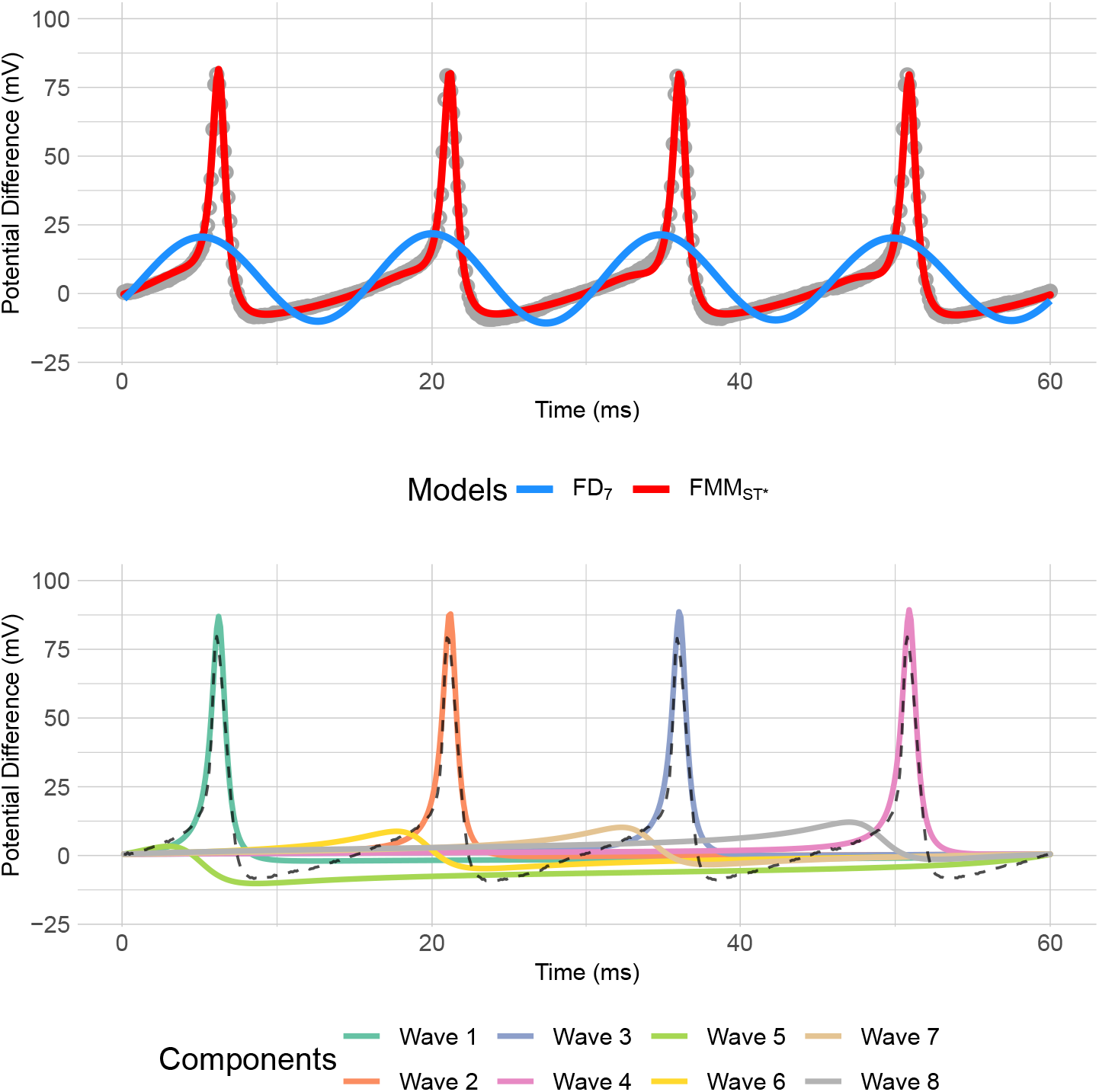
A neuronal AP from the SGAMP database (axon 1, trial 18, second short square stimulus) along with the fitted signals using an FMM_ST*_ signal (red) and an *FD*_7_ (blue). The components of the fitted FMM_ST*_ model are illustrated at the bottom.

Finally, a principal component analysis has been performed using the basic set of FMM parameters from the HH experiments. A plane for the first two principal components is given in Fig. 6, where each point represents an HH simulated signal. In that plot, the values corresponding to SGAMP data have also been plotted as they are located far from the main cloud of HH points.

**Fig 6.**
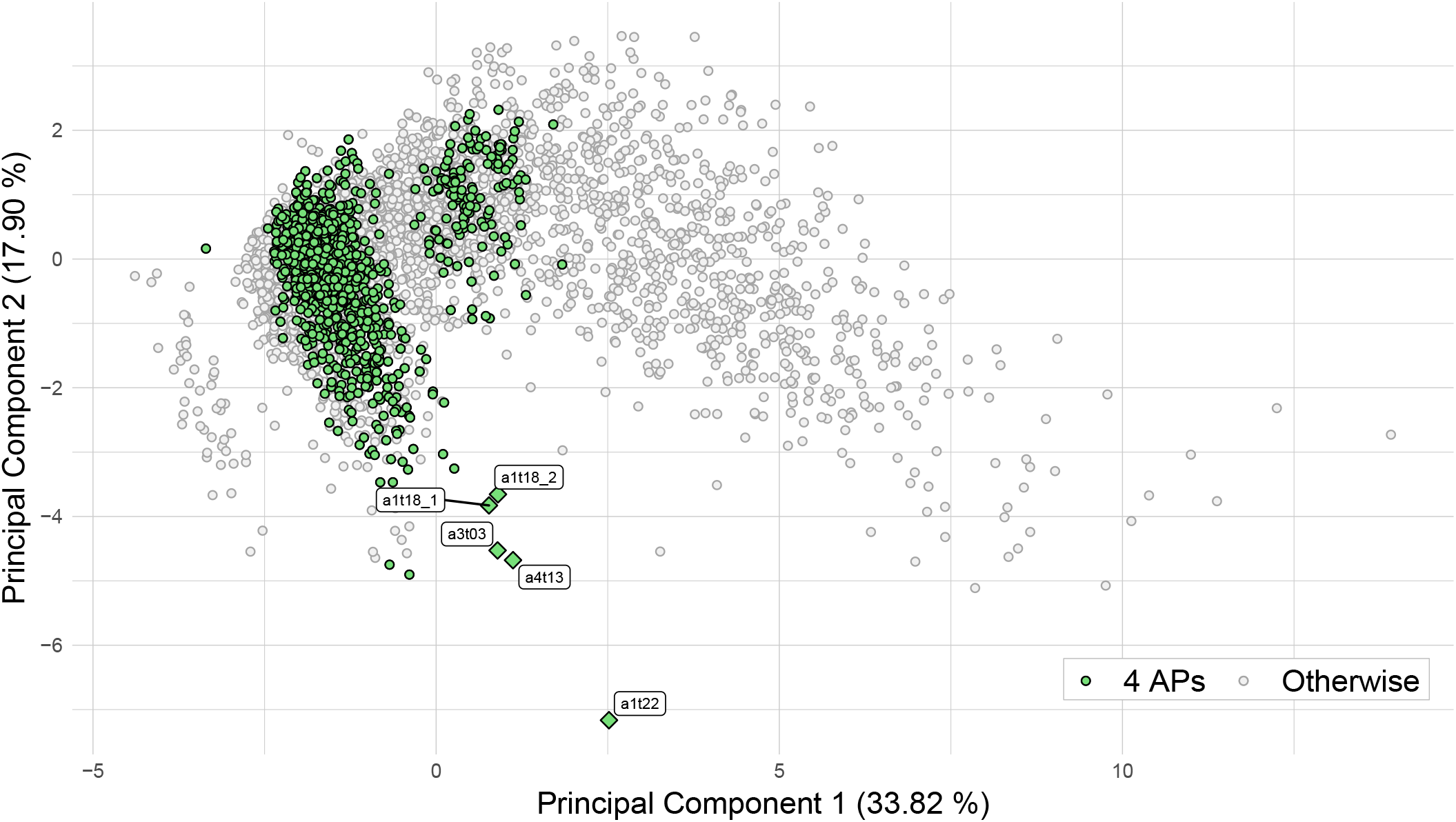
Plot of the first two principal components of *τ^A^* + *τ^B^* of the HH experiments. Experiments with 4 APs observed have been highlighted. The projections of the SGAMP experiments have been added to the plot with the corresponding label.

This latter fact, and the results shown above the percentage of variance explained by the Dominant Component, indicate that the model underlying SGAMP signals is not an HH but a simpler one, such as FitzHugh-Nagumo (see [9]).

## Discussion

In this paper, the FMM_ST_ model has been presented, and its potential to predict simulated and real AP signals has been proved. Our analysis shows that some differences in shape exist between HH and SGAMP signals, even though it has often been assumed that the HH model is adequate to represent the SGAMP data in the literature. Therefore, a simpler model may be more suitable to represent SGAMP signals. Moreover, the accurate prediction of HH and SGAMP signals also provides evidences that other neuronal dynamics models could also be represented using the FMM approach. Differences between models can be articulated through the differences in the FMM parameter configurations.

Furthermore, the paper shows the potential of the model to describe such neuronal features as 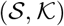. In particular, the flexibility of the FMM parameters to describe different AP waveforms would contribute to solving problems where the AP characterization is needed. Specifically, it is relevant in Spike Sorting, which is the process in which spikes are grouped into clusters based on the similarity of their shapes. The resulting clusters are assumed to correspond to different neurons. Therefore, Spike Sorting is the first step in many data analyses, being one of the most critical data analysis problems in neurophysiology and has received a lot of attention in the literature. Some interesting references are [24], [43] and [44], among others.

The proposed approach is also flexible concerning the segment length. The signal can be analyzed, either for individual AP segments, which implies spike cutting as in other models, such as [13], or by analyzing a more extended length of signal, most likely with several APs, as is done in this work.

Two different lines of work could be defined for the future. On the one hand, from a theoretical perspective, a first question to solve is the implementation of restrictions with the form 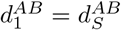; *S* = 2*,…, s*. They are of interest to analyze signals with equal distances between components in the APs, because reducing the number of parameters to be estimated is important when large or noisy Spike Trains are analyzed. On the other hand, from an applied perspective, many other AP real signals must be analyzed, and the questions of spike classification and clustering may be addressed.

Particularly, it would be of interest to consider the FMM parameters in the problem of neuronal type classification, as they may be useful to characterize the electrophysiological mechanisms, morphological features, and genetic profiles. Some works in this line are [2], [13] and [45]. The FMM approach’s potential is difficult to calibrate as many aspects remain to be researched and exploited.

Finally, a limitation of our study is that the input stimulus’s timing and shape have been fixed. The influence of the stimulus type could be analyzed using the FMM approach. The *α* parameters are related to the firing times, and the shape parameters could be useful to detect circumstances where the shape of the APs is independent of the stimulus, as [4] suggests, and circumstances where changes happen, such as the observation of incomplete spikes. However, the question is tricky and deserves further research.

